# An engineered biofactory for efficient production of diverse recombinant superoxide dismutase isozymes loaded with specific metal ions for biochemical characterization

**DOI:** 10.64898/2026.07.08.737244

**Authors:** Mariam Esmaeeli, Agnieszka Kołpa, Rafał Mazgaj, Justyna Pełczyńska, Diana Galea, Jan Gawor, Agata Malinowska, Agnieszka Szczypiorowska, Thomas E. Kehl-Fie, Kevin J. Waldron

## Abstract

**Background:** Biochemical, biophysical and structural characterisation of isozymes from the ubiquitous family of iron- or manganese-dependent superoxide dismutases (SodFMs) requires the purification of high-quality preparations of recombinant enzymes. Determination of their key biochemical parameter, their catalytic metal-preference, requires the comparison of the catalytic turnover of samples loaded exclusively with iron versus samples loaded exclusively with manganese. Both of these aims are inhibited by the potential contamination of recombinant preparations of SodFMs, prepared by heterologous overexpression inside *Escherichia coli* cells, by even low levels of endogenous SodFMs from the host, both of which show very high turnover with either manganese (*E. coli* MnSOD) or iron (FeSOD). To overcome this problem, we created a strain of *E. coli* lacking the endogenous SodFMs. Here, we characterised this *E. coli* BL21 (DE3) Δ*sodA*Δ*sodB* strain, determining the physiological effects of SodFM deletion and demonstrating its utility for producing recombinant SodFMs for *in vitro* characterisation and use.

**Results:** Genomic analysis verified the targeted gene deletions, without off-target effects. Growth, expression, elemental analysis, and proteomic data confirmed a lack of physiological defects of the strain except for a known inability to grow on glucose, which is overcome by heterologous SodFM expression. We demonstrate the utility of the strain for the efficient production of diverse recombinant SodFMs, including highly divergent, understudied isozymes, including the ability to precisely control the metal-loading of the heterologously expressed protein.

**Conclusions:** The *E. coli* strain described herein is a useful microbial cell factory for production of recombinant SodFMs, which should find widespread utility as expression host of choice, enabling more efficient production of protein for studies of the biochemical, biophysical and structural properties of this remarkable family of metalloenzymes.

## Background

Since the first expression of a recombinant protein inside cells of *Escherichia coli* was reported in 1977 [1] and the first recombinant therapeutic, human insulin, in 1979 [2], demand for recombinant protein production for academic study and for commercial, medical or industrial applications has grown dramatically. Although alternatives are available, *E. coli* remains the cell factory of choice for producing recombinant proteins [3,4]. It offers several advantages for recombinant protein synthesis. It has been extensively studied, with well-characterised physiology and genetics and easy-to-use tools available [5–10]. These include inducible expression vectors [6,7,11] and the host strain, BL21 (DE3). This strain lacks Lon and OmpT proteases, limiting degradation of foreign protein, and incorporates RNA polymerase from T7 bacteriophage in its genome [12], enabling it to express transcripts from the strong T7 promoter [5].

Derivatives of *E. coli* BL21 (DE3) have been constructed to improve its production of proteins for specific applications. Strains were created that are optimised for production of proteins containing a poly-histidine tag to facilitate their purification. These strains have mutations in five endogenous *E. coli* genes whose products commonly contaminate protein preparations after purification by immobilised metal affinity chromatography (IMAC), reducing their affinity for IMAC or enabling a pre-clearance step using chitin [13]. Other modified BL21 strains eliminate contamination of recombinant protein preparations by the chaperone DnaK [14], improve expression of outer membrane proteins [15], facilitate site-specific incorporation of non-natural or isotopically labelled amino acids [16,17], produce glycosylated eukaryotic proteins [18], or improve production of specific metabolites such as N-acetyl-D-neuraminic acid (Neu5Ac) [19], lycopene [8], or phenylpyruvic acid [20].

For the last decade, we have been studying the iron- or manganese-dependent superoxide dismutase (SOD) family of metalloenzymes (SodFM) [21–27]. SODs are the primary cellular defence against superoxide radicals, key antioxidant enzymes that maintain redox balance, and in higher eukaryotes have been implicated in signalling, immune function and disease [28]. The SodFMs are the most common and most widely distributed class of SODs, present in diverse organisms including some obligate anaerobes [21]. SodFMs exhibit a spectrum of metal specificities, ranging from highly manganese-preferring to highly iron-preferring enzymes [21,29,30]. Between these extremes are isozymes that can catalyse their reaction with either metal cofactor, termed cambialism [21,24,31], a rare biochemical property among redox metalloenzymes [32]. All SodFMs share identical folds and remarkably similar active site architectures [21,28,33,34], leading to questions about how different isozymes control their distinct catalytic metal-preferences [22,23,28,35]. The detailed biochemical, biophysical and structural characterisation of SodFMs needed to answer such questions requires purified preparations of recombinant enzymes, free of contamination by the endogenous SodFMs that are expressed by the host cell. *E. coli* possesses two SodFMs, strongly manganese-preferring *Ec*SodA and iron-preferring *Ec*SodB [36–38], which are highly active enzymes with their cognate metal cofactor [21,29,30]. This makes even very low levels of their contamination in recombinant preparations of heterologous SodFMs highly problematic for quantifying the catalytic turnover of an isozyme with each metal, especially as highly metal-preferring isozymes can have very low catalytic efficiency when loaded with the ‘wrong’ metal cofactor [21,23,29,30].

To overcome this bottleneck, we created a strain of *E. coli* BL21 (DE3) in which both genes encoding SodFMs were deleted from the genome [21]. The strain grows normally in aerobic culture when used for recombinant protein expression [21,39]. The BL21 (DE3) Δ*sodA*Δ*sodB* strain enables large-scale preparation of recombinant SodFMs, free of contamination by native *E. coli* SodFMs, and with specific metal-loading despite the BL21 parental strain being known to have defects in metal homeostasis [40]. Using this strain, we developed medium throughput in-gel assays to assess the catalytic metal-preference of SodFM isozymes, which enabled a study of the evolution of the SodFM family [21]. Here, we have characterised this *E. coli* mutant strain, indicated its lack of physiological growth defects under standard culture conditions, and demonstrated its usefulness as an expression host for the production of diverse SodFM enzymes for downstream studies.

## Methods

### Bacterial strains, constructs and culture media

The *E. coli* BL21 (DE3) strain used was derived from a batch of competent cells purchased from Thermo. The construction of the strain lacking both endogenous SodFM-encoding genes within this BL21 strain was described previously [21]. Competent cells of each strain, wild type (WT) and Δ*sodA*Δ*sodB*, were produced using a standard Mg/Ca method and transformed with a pET22b construct for expression of a heterologous SodFM isozyme from our library, also previously described [21], or with pET22b empty vector. Cells were cultured in LB medium or in M9 medium containing selected supplements, with 100 µg/mL ampicillin added to select for the pET22b plasmid, at 37°C with 140 rpm orbital shaking. Growth analyses (in biological triplicate) were performed in 96-well microtitre plates by continuously monitoring OD_600nm_ at 37°C with 140 rpm orbital shaking using a Synergy H1 microplate reader (BioTek). Glucose or glycerol (0.2% w/v) as carbon source, casamino acids (0.4% w/v) or 100 µM MnCl_2_, were added to 200 µL M9 minimal medium. Cell samples for proteomics were cultured (100 mL LB) in batch flasks. Cell samples for testing of recombinant protein expression and activity were cultured (50 mL in M9 medium supplemented with metals as previously described [21] to control metal-loading)

### DNA isolation, sequencing and genomic analysis

Short-read bacterial genome sequencing was performed using the MiSeq instrument (Illumina Inc., San Diego, CA, USA) at Genomed S.A. (Warsaw, Poland). Genomic DNA was purified using the PureLink Genomic DNA Mini Kit (Thermo Fisher Scientific, Waltham, MA, USA). DNA quality control was performed by measuring the absorbance at 260/230 using Nanodrop Spectrophotometer (Thermo Scientific) and DNA integrity was analyzed by 0.5% agarose gel electrophoresis. DNA libraries were constructed using an NEB Ultra II FS kit (NEB, Ipswich, MA, USA), followed by paired-end 300 basepair sequencing (aiming for at least 50x genome coverage). Sequence quality metrics were assessed using FASTQC v.0.12.1 (http://www.bioinformatics.babraham.ac.uk/projects/fastqc/, accessed on 31 January 2025). Raw sequencing reads were trimmed for quality and residual library adaptors were removed using fastp v.0.23.4 (accessed on 31 January 2025) [41]. Cleaned sequencing reads were then processed by breseq v.0.39.0 pipeline [42] to map reads to the reference genome (GenBank accession number: CP001509) and identify variants. Illumina reads for DeltaBL21 mutant were additionally assembled into contigs using the Unicycler v.0.4.8 pipeline (accessed on 31 January 2025) [43]. De novo assembled contigs were annotated using DFAST v.1.3.0 [44]. GenBank annotation files for the DeltaBL21 mutant strain and the *E. coli* BL21 reference genome were used for sequence alignment and visualization using Clinker v0.0.31 [45].

### Proteomic analyses

Parallel cultures of *E. coli* BL21 (DE3) WT and the Δ*sodA*Δ*sodB* strain, each transformed with pET22b-*Sa*SodM, were inoculated into LB. The initial batch of samples was collected after 2 h 40 min at mid-log phase. The cultures were then induced with 100 µM IPTG and the second batch of samples was collected after 4 h induction. For sample collection, 1 mL culture was removed, cells harvested (5,000 *g*, 10 min, 4°C), then washed twice in 1 mL wash buffer (20 mM Tris, pH7.5, 150 mM NaCl) with centrifugation (6,000 *g*, 10 min, 4°C), and pellets stored at −80°C. This procedure was repeated for 5 independent biological replicates on different days.

For lysis, pellets were thawed on ice and resuspended in 300 µL of lysis buffer (100 mM NH_4_HCO_3_ with 2% SDS), then samples sonicated in an ice bath (10 cycles: 1 min on, 2 min off). Samples were heated at 95°C for 15 min and centrifuged (12,000 *g*, 20 min, 4°C), the supernatant removed and transferred to fresh tubes, frozen in liquid nitrogen and kept at −80°C.

After thawing, the protein concentration in each sample was estimated using a bicinchoninic acid (BCA) assay using a Pierce™ BCA Protein Assay Kit (Thermo Scientific) for mass spectrometry (MS) analysis at the Mass Spectrometry Laboratory at the Institute of Biochemistry and Biophysics, Polish Academy of Sciences (PAS). The samples (35 µg protein) were mixed with 1% SDS in 100 mM NH_4_HCO_3_, cysteines reduced by 1 h incubation with 5 mM tris(2-carboxyethyl)phosphine (TCEP) at 60°C followed by 10 min incubation at room temperature with 20 mM methylmethanethiosulfonate (MMTS), then processed using single-pot solid-phase-enhanced sample preparation (SP3). Magnetic beads were prepared by combining equal parts of Sera-Mag carboxyl hydrophilic and hydrophobic particles (09-981-121 and 09-981-123, GE Healthcare). The bead mix was washed three times with MS-grade water and resuspended at a working concentration in 85% acetonitrile (ACN). The bead mix was added to the samples, which were suspended in 0.1% formic acid in ACN. Using the magnet beads, the samples bound to the bead mix were separated, washed in 100% isopropylalcohol, then washed three times in 85% ethanol, then twice in 100% ACN. Digestion was performed by incubation at 37°C overnight with 1 µg of trypsin (Promega) in 100 mM NH_4_HCO_3_, then peptides were eluted from the beads using 1% dimethyl sulfoxide (DMSO) in MS-grade water. Peptides were dried in SpeedVac and resuspended in 60 µL of 0.1% formic acid in MS-grade water with sonication.

Peptide concentrations were measured using Colorimetric Peptide Assay (Thermo Scientific). Peptide mixtures from each sample (2 µg) were analysed using an LC-MS system consisting of an Evosep One (Evosep Biosystems) coupled with an Orbitrap Exploris 480 mass spectrometer (Thermo Scientific). The raw data were submitted to MaxQuant/Andromeda (version 2.5) for peptide and protein identification, and calculation of quantitative values. For peptide and protein identification, a UniProt-derived reference database for *E. coli* (strain B/BL21-DE3, 4,156 sequences, version2024_06) was used. The database was supplemented with the sequence of the *S. aureus* SodM protein (Q2G261) and a database of common mass spectrometry (MS) contaminants. Further analysis was performed using Perseus (version1.6.15).

### Recombinant protein expression and analyses

For heterologous expression of SodFM enzymes, *E. coli* BL21(DE3) wild type and the isogenic Δ*sodA*Δ*sodB* strains were transformed with a pET22b(+) plasmid construct encoding one of the following SodFM enzymes: SodFM1 from *Staphylococcus aureus* (*Sa*SodA), SodFM2 from *Neisseria gonorrhoeae*, SodFM3 from *Mycobacterium abscessus*, SodFM4 from CPR *Parkubacteria*, or SodFM5 from *Homo sapiens* [21].

Overnight cultures were grown in M9 minimal media supplemented with 50 µg/mL ampicillin at 37°C with 140 rpm shaking, then diluted 1:100 into fresh M9 medium with 50 µg/mL ampicillin. Cultures were grown at 37°C until mid-log phase (OD_600nm_ ∼0.5), induced with 100 µM IPTG and supplemented with 200 µM of the appropriate metal (MnCl_2_ or FeSO_4_). Induced cultures were incubated at 37°C for 16 h. Cells were harvested by centrifugation (4000 *g,* 15 min, 4°C), pellets washed twice with 20 mM Tris pH 7.5 150 mM NaCl 10 mM EDTA and stored at −20°C.

Whole-cell extracts were prepared by resuspending cell pellets in wash buffer, 100 mg/mL lysozyme added followed by incubation for 30 min with gentle rocking, followed by 4 freeze-thaw cycles with liquid nitrogen. Cell lysates were clarified by centrifugation (20,000 *g*, 20 min, 4°C), and the soluble fraction was used for further analyses.

Expression was confirmed by SDS-PAGE, total protein concentration was determined by BCA assay using a commercial kit (ThermoFisher).

### In-gel activity staining

Enzymatic activity of the recombinant SodFM proteins was measured in the clarified lysates using in-gel activity assay [21], and activities were normalized to total protein content. Protein samples, prepared in native gel protein loading buffer, were resolved on native 12% (v/v) polyacrylamide gels lacking SDS. Metal-specific activity was assessed by incubation of one of two identical gels for 30 min in solution of 0.3% (v/v) H_2_O_2_ at 4°C prior to activity staining. Gels were subsequently washed and incubated with a staining solution of 0.25 mM nitro blue tetrazolium, 0.03 mM riboflavin, 0.1 mM TEMED (N,N,Ń,Ń-tetramethylethylenediamine) for 30 min at 4°C. Gels were then washed in water and placed on a white light transilluminator for development of the in-gel activity staining. The gels were exposed to equal white light for standardised time. Images of the gels were acquired with a Bio-Rad ChemiDoc System.

### Inductively coupled plasma optical emission spectrometry elemental analysis

For ICP-OES analysis, 500 µL of ultrapure nitric acid (Merck) was added to the cell pellet, and the mixture was digested at 80°C for 2 h. Digested samples were diluted with 1% (v/v) nitric acid prior to measuring the metal content. Elemental analysis was performed on an iCAP Pro ICP-OES instrument (Thermo-Fisher Scientific) run in aqueous-axial-iFR mode, equipped with an iSC-65 autosampler and a Torch Duo (Slot) Rev 02 argon plasma torch set, at 1,250 W RF power and using Qtegra software. An external calibration curve was recorded with ICP-single-element standards mix in 1% (v/v) nitic acid. The sample was introduced via a peristaltic pump for analysis. The wavelengths for S (182.034 mm), Zn (206.200 nm), Mn (257.610 nm), Fe (259.940 nm), Mg (285.213 mm) were selected to minimise overlapping emissions. The results were calculated from the ppb data, normalized to the S content.

### Gene expression analysis by quantitative PCR (pQCR)

*E. coli* BL21 (DE3) WT and the Δ*sodA*Δ*sodB* strain, each transformed with either pET22-SodM or the empty vector, were inoculated into LB medium. Overnight cultures were diluted 1:100 and cultivated at 37 °C with shaking at 140 rpm. When the cultures reached an OD_600nm_ of 0.5, protein expression was induced with 100 µM IPTG, followed by incubation for an additional 4 h. Aliquots (1 mL) of each culture were harvested by centrifugation (5,000 × g, 10 min, 4 °C).

Total RNA was isolated using a PureLink RNA Mini Kit (Invitrogen). Genomic DNA was removed by on-column DNase digestion with the PureLink DNase Set (Invitrogen). For real-time quantitative PCR (RT-qPCR), 2 µg of total RNA was reverse-transcribed using the Maxima First Strand cDNA Synthesis Kit (Thermo Fisher Scientific, Waltham, MA, USA) according to the manufacturer’s instructions. The obtained cDNA was diluted 200-fold. RT-qPCR was performed on a LightCycler 480 system (Roche, Basel, Switzerland) using the Luminaris Color HiGreen qPCR Master Mix (Thermo Fisher Scientific, Waltham, MA, USA). The cycling conditions consisted of an initial denaturation at 95 °C for 6 min, followed by 40 cycles of denaturation at 95 °C for 5 s and annealing at 60 °C for 30 s. The *rrsA* and *hcaT* genes, encoding 16S ribosomal RNA and a putative 3-phenylpropionic acid transporter, respectively, were used as reference genes. Primers used for qRT-PCR are listed in Supp. Table S1. Each sample was measured in triplicate, and four biological replicates were used. Relative gene expression levels were calculated using the ΔΔCt method [46]. Statistical significance was assessed using Student’s t-test.

## Results

### The genome of the BL21-derived host strain shows no off-target mutations

The *E. coli* BL21 strain is known to be vulnerable to genetic mutations, showing an increased mutation rate relative to other Type B *E. coli* [47]. The Lambda Red method for genetic modification of the *E. coli* genome has also been suggested to be vulnerable to secondary mutations due to the use of low and high temperature selection steps [48,49]. Further, deleting genes encoding superoxide dismutases, which are involved in cellular defence against exogenous oxidative stress, also makes the host cell vulnerable to secondary mutations [50]. For these reasons, we began our study of the BL21 Δ*sodA*Δ*sodB* strain by sequencing its genome to identify any mutations that were present in addition to the targeted deletions.

We used Illumina sequencing to compare the genome sequences of both the Δ*sodA*Δ*sodB* strain and the parental BL21 (DE3) strain from which it was made (herein ‘wild type’, WT) to the reference genome for *E. coli* BL21 (DE3) [51]. The WT strain showed no deviations from that of the reference genome. In the Δ*sodA*Δ*sodB* strain, as expected, clean deletions of the *sodA* and *sodB* genes were observed, with the genes replaced by scar sequences (Fig. 1A, 1B), a remnant of the flippase recombination event that excised a selectable marker during the mutagenesis protocol [48]. Consistent with this, we detected three bands of abundant SOD activity in crude lysates prepared from BL21 (DE3) cells using an in-gel activity assay, corresponding to homodimers of *Ec*SodA and *Ec*SodB plus a heterodimeric form, as previously observed [52]. All three bands were absent from lysates prepared from the BL21 Δ*sodA*Δ*sodB* strain (Fig. 1C).

**Figure 1:**
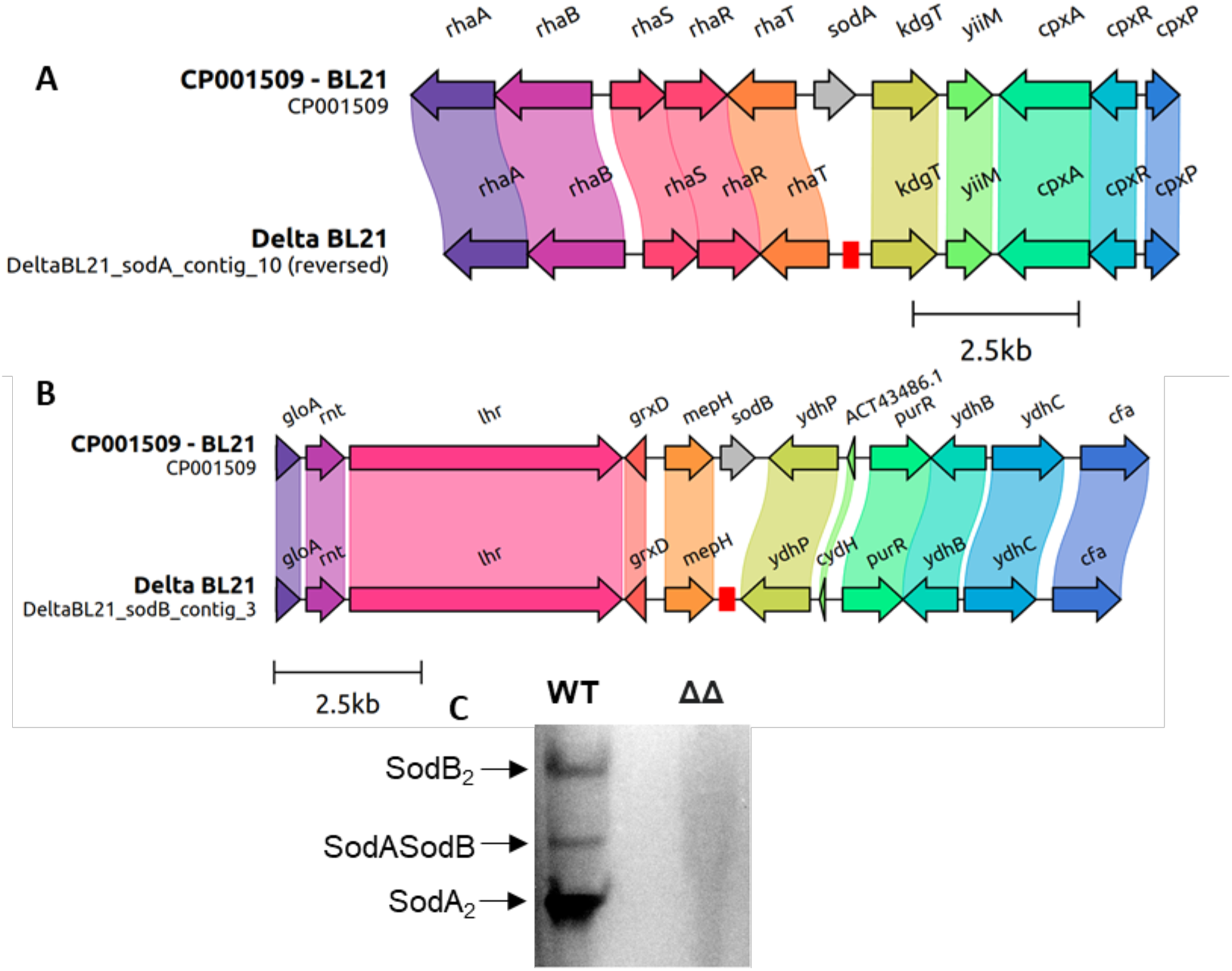
Genome and in-gel activity assay confirms deletion of SOD genes. A-B. Schematic representations of the alignment of the (A) *sodA* and (B) *sodB* gene regions between the *E. coli* BL21 (DE3) reference genome (CP001509) and the BL21 Δ*sodA*Δ*sodB* strain (Delta BL21). Scar sequences in the mutant strain are highlighted with red boxes. C. Equal amounts (10 µg) of crude soluble extracts from *E. coli* BL21 (DE3) wild type (WT) and from the *E. coli* BL21 Δ*sodA*Δ*sodB* strain (ΔΔ) were resolved by native PAGE on a 15% acrylamide gel, and SOD activity detected with NBT/riboflavin straining.

Aside from the two targeted deletions, the only other mutation we observed in the Δ*sodA*Δ*sodB* strain was a single SNP (Table 1). This was an A◊G mutation within the region between the *sulA* and *sxy* genes. Inspection of that region suggested that this SNP was unlikely to impact on the transcription of either of the neighbouring genes [53]. Thus, the construction of the expression host strain from wild type BL21 had been remarkably faithful, despite the potential risks posed by this method of manipulation, the parental genotype, and the potential effects of the predicted phenotype that might be caused by the deletions.

**Table 1:** Mutations unique to mutant (ΔΔ) and wild type (WT).

| Mutations present in $\Delta\Delta$ and not present in WT | | | | |
| --- | --- | --- | --- | --- |
| Position | Mutation | Annotation | Gene | Description |
| 1,026,050 | A→G | Intergenic<br>(-42/-177) | *sulA* ← / → *sxy* | SOS cell division inhibitor/CRP-S-dependent promoter expression factor |
| 1,680,577 | 558 bp→84 bp | Coding<br>(4-561/582 nt) | *sodB* → | superoxide dismutase, Fe |
| 4,008,735 | 597 bp→84 bp | Coding<br>(4-600/621 nt) | *sodA* → | superoxide dismutase, Mn |
| Mutations present in WT and not present in $\Delta\Delta$ | | | | |
| 1,136,454 | C→T | A29V<br>(GCC→GTC) | *flgF* → | flagellar component of cell-proximal portion of basal-body rod |
| 2,006,172 | G→A | A338V<br>(GCC→GTC) | *vioA* ← | VioA, involved in dTDP-N-acetylviosamine synthesis |

List of mutations identified by breseq pipeline after mapping sequencing data to the *E. coli* BL21 (DE3) $\Delta\text{sodA}\Delta\text{sodB}$ strain and the wild-type reference genome, respectively.

### Growth of the E. coli BL21 ΔsodAΔsodB strain was indistinguishable from wild type

Previous reports showed that a Δ*sodA*Δ*sodB* strain of Type A *E. coli*, the K-12 strain MC4100, exhibited a severe growth defect under certain growth conditions. Specifically, this strain was unable to grow in media containing glucose as the sole carbon source, but was able to grow on alternative carbon sources such as amino acids [39]. Subsequent studies have indicated that superoxide stress strongly affects enzymes containing exposed iron-sulphur clusters, including enzymes in central carbon metabolism and the biosynthesis of branched-chain amino acids [54–56]. Therefore, this failure of the *E. coli* MC4100 Δ*sodA*Δ*sodB* strain to grow in glucose was proposed to be a result of a superoxide-induced metabolic block [39,55].

Consistent with this, we observed only a minor growth defect of the Δ*sodA*Δ*sodB* strain, carrying an empty vector, when cultured in LB medium, but a severe growth defect of the same strain in an M9 minimal medium supplemented with glucose or with glycerol as sole carbon source (Fig. 2A). Growth of the double knockout strain was significantly better when the medium was supplemented with amino acids. Growth was also improved by supplementation of the medium with excess manganese (Fig. 2A), which can suppress oxidative stress in *E. coli* [55,57,58]. In all media, provision of a heterologous SodFM abolished the phenotype. Complementation of the Δ*sodA*Δ*sodB* strain with a pET22 vector carrying the *S. aureus* SodM (*Sa*SodM) [21] resulted in growth similar to the WT strain (Fig. 2A). We selected this isozyme because it is cambialistic, i.e. can catalyse its dismutation reaction equally well using either a manganese or iron cofactor [23,24], thus ensuring molecules of *Sa*SodM would be catalytically active inside the *E. coli* cells regardless of which of these metal ions they acquired. Together, these data demonstrate that the *E. coli* BL21 Δ*sodA*Δ*sodB* strain exhibits growth phenotypes that are analogous to those previously observed in an *E. coli* K12 strain lacking its SodFMs [39], and these phenotypes can be attributed to loss of SOD activity.

**Figure 2:**
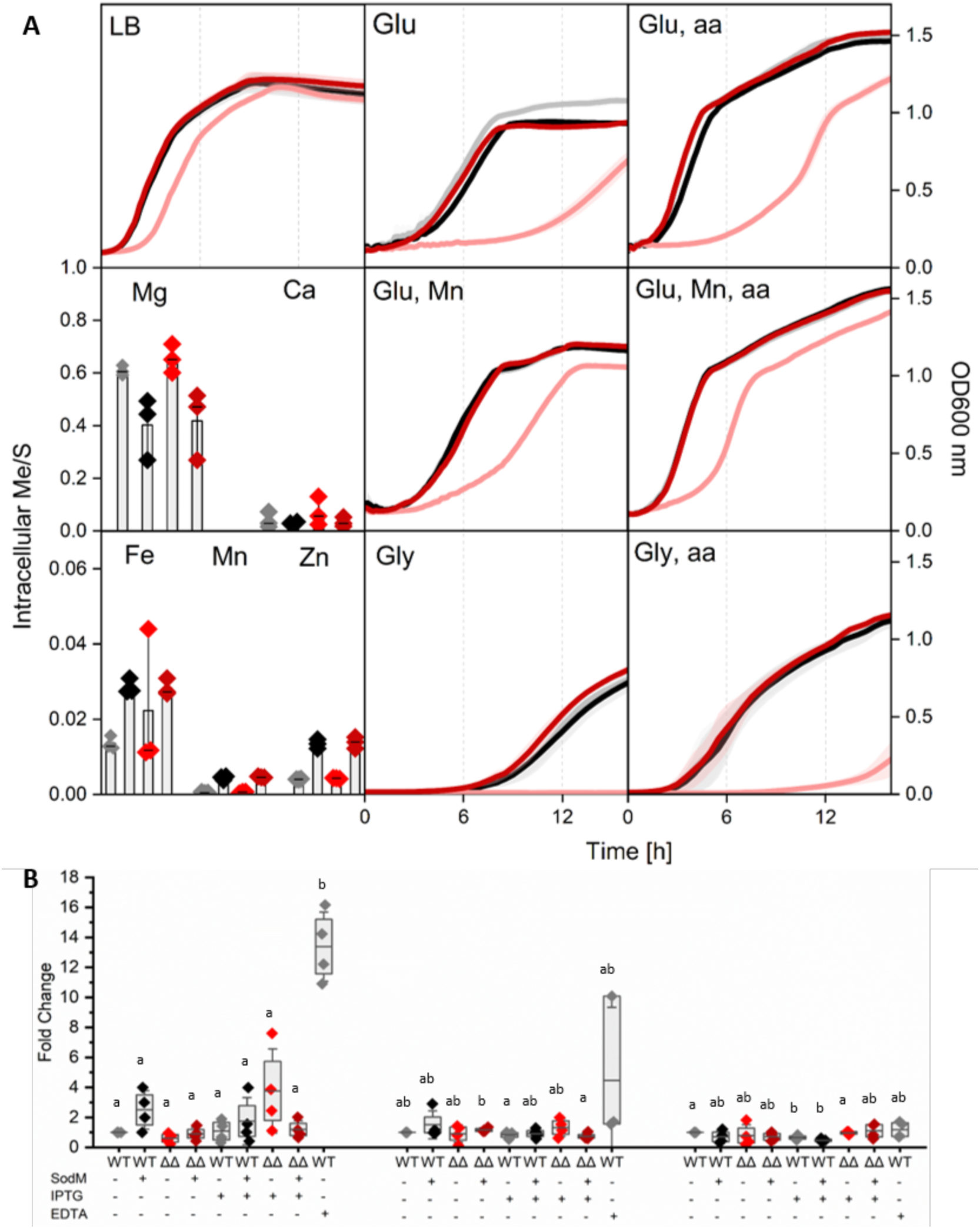
*E. coli* Δ*sodA*Δ*sodB* strain shows expected growth phenotype but no defects in metal homeostasis. A. Growth and total metal accumulation phenotypes of the *E. coli* Δ*sodA*Δ*sodB* strain. Growth of *E. coli* BL21 (DE3) WT (grey) and the Δ*sodA*Δ*sodB* strain (red) strains, carrying either a pET22 empty vector (lighter colours) or with the pET22-*Sa*SodM construct (darker colours), were assessed either in a rich LB medium, or in M9 minimal medium supplemented with glucose (Glu), glycerol (Gly), casamino acids (aa), or with 100 µM MnCl_2_ (Mn). Growth curves, in biological triplicate with mean values shown, were assessed through continuous monitoring of OD_600nm_ in a microtitre platereader at 37°C with 140 rpm orbital shaking. Samples (1 mL) of cells of the same strains were harvested at mid-log phase (OD_600nm_ ∼0.5) during growth in LB medium, digested in nitric acid, and their elemental composition was quantified by ICP-OES for magnesium (Mg), calcium (Ca), iron (Fe), manganese (Mn) or zinc (Zn). B. Total RNA was extracted from samples (1 mL) of cells from LB cultures of the same strains. Cells were cultured to mid-log (OD_600nm_ ∼0.5) then expression of *Sa*SodM was induced by addition of 100 µM IPTG in one set of cultures and left uninduced in another set, followed by 4 h further incubation at 37°C with 140 rpm shaking. A control culture was exposed to 10 mM EDTA in the medium for 30 minutes. After RNA purification and genomic DNA digestion, RNA samples (2 µg) were reverse-transcribed, and resulting cDNA used for qPCR analyses. The abundance of transcripts for *mntH*, *feoA* and *fecA* were quantified, while the *rrsA* and *hcaT* genes were used as reference genes. Each sample was measured in technical triplicate, across four biological replicates, and relative gene expression calculated using the ΔΔCt method with mean and SD shown by box-and-whiskers, and the individual values illustrated. Statistical significance was assessed using Student’s *t*-test, indicated by the letters.

We next investigated metal homeostasis in the BL21 Δ*sodA*Δ*sodB* strain to assess its suitability for overproduction of a metalloenzyme. We compared the metal content of cells of the Δ*sodA*Δ*sodB* strain relative to its parental WT cells using inductively coupled plasma optical emission spectrometry (ICP-OES) (Fig. 2A). The data demonstrated no significant differences in the cellular Mg, Ca, Fe, Zn or Mn content between the strains, transformed with either the SodM-expression plasmid or an empty pET22 vector. We also assessed the transcription of known manganese and iron uptake genes in *E. coli* [59–61] by reverse-transcription quantitative polymerase chain reaction (RT-qPCR) to assess whether the Δ*sodA*Δ*sodB* strain displays a metal-deprived phenotype. We observed no significant change in the expression of the gene encoding the manganese-importer MntH [58,62] in the Δ*sodA*Δ*sodB* strain, nor of either of genes that encode the primary iron-importers under standard culture conditions, FeoA (which transports ferrous ions) or FecA (which transports ferric citrate) [60,63] (Fig. 2B). These data indicate that the Δ*sodA*Δ*sodB* strain does not suffer from an inherent defect in metal homeostasis.

### Proteomic analyses indicate few differences between cells of E. coli BL21 WT and the ΔsodAΔsodB strain under recombinant protein expression conditions

We next sought to investigate the cellular physiology in the *E. coli* BL21 Δ*sodA*Δ*sodB* strain and its usefulness as a microbial expression host for the production of heterologous SodFM isozymes. We used proteomic analyses to measure changes in protein expression in this strain relative to its WT parent during overexpression of the heterologous proteins. We quantified proteins from each strain, transformed with the pET22 construct containing the *sodM* gene from *S. aureus*, in LB medium with (100 µM IPTG) or without induction of expression of the heterologous gene, to determine how the cellular proteome was affected by deletion of *sodA* and *sodB* and by the induction of the heterologous SOD (Supp. Fig. S1).

When we compared proteomes of the wild type to the Δ*sodA*Δ*sodB* strain prior to induction of the heterologous protein, we identified just 50 proteins whose abundance had significantly changed, 42 that were up- and 8 that were down-regulated (Fig. 3A). This small number of changes indicated an absence of serious metabolic or physiological problems in the Δ*sodA*Δ*sodB* strain when carrying an expression construct encoding a functional, heterologous SodFM, consistent with leaky expression from the pET system being sufficient to provide the cell with protection against endogenous oxidative stress. Indeed, we detected the presence of *S. aureus* SodM in both proteomes, and its abundance was found not to be statistically significantly different when comparing between the extracts from WT vs the Δ*sodA*Δ*sodB* cells (Fig. 3A).

**Figure 3:**
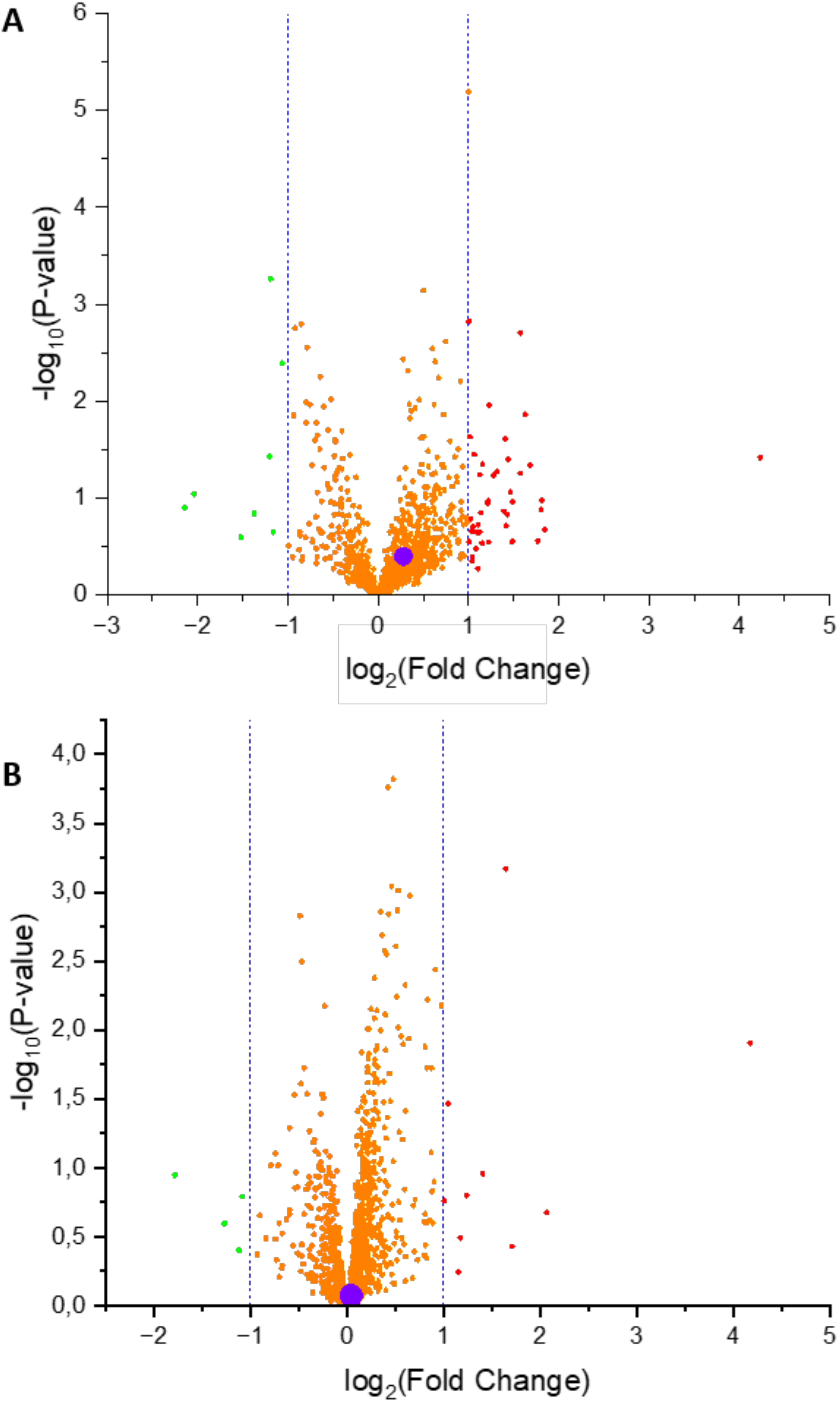
Proteomic comparison of wild type and mutant BL21 cells shows production strain lacks physiological defects. Proteins were extracted from aliquots (1 mL) of cultured *E. coli* BL21 (DE3) wild type and from the *E. coli* BL21 Δ*sodA*Δ*sodB* strain, each carrying the pET22b-*Sa*SodM construct, before and after induction (100 µM IPTG) of expression of the heterologous *Sa*SodM protein. Cells were lysed in SDS buffer by sonication, equal amounts of protein (35 µg) digested with trypsin, and resulting peptides were purified. Peptide mixtures (2 µg) analysed by LC-MS. Peptides were assigned to proteins with MaxQuant using an E. coli reference database, and then changes in protein abundance analysed with Perseus. The resulting volcano plots are presented for comparison between WT and ΔΔ strains (A) without added IPTG inducer and (B) with induction of the heterologous protein. The red circles represent proteins that were up-regulated (>1 log_2_), the green circles represent the proteins that were down-regulated (<1 log_2_), and all proteins found to show no change in abundance are represented by orange circles. The large blue circle represents the heterologously expressed *S. aureus* SodM protein. The dotted blue lines represent the log2 fold-change cutoff used.

Likewise, when we compared the proteome of the same cells after induction of the heterologous *Sa*SodM, we identified even fewer proteins whose abundance had significantly changed, 10 that were up- and 4 that were down-regulated (Fig. 3B). This indicated the physiology of the Δ*sodA*Δ*sodB* strain is highly similar to that of WT BL21 cells, when over-expressing *Sa*SodM. Importantly, the abundance of *S. aureus* SodM was unchanged in the two proteomes, showing that the Δ*sodA*Δ*sodB* cells are as efficient for production of *Sa*SodM as the WT BL21 cells (Fig. 3B).

As expected, we observed far more proteomic changes within each strain when we compared induced to uninduced samples (Supp. Fig. S1). We detected 125 proteins whose abundance was significantly changed (89 increased, 36 decreased) by induction of the recombinant protein within wild type cells (Supp. Fig. S2), reflecting the physiological effects on *E. coli* BL21 (DE3) cells caused by the overexpression of the heterologous *Sa*SodM isozyme. We also observed 223 proteins whose abundance was significantly changed (163 increased, 60 decreased) by induction of the recombinant protein within Δ*sodA*Δ*sodB* cells (Supp. Fig. S3). The latter included changes in sulfur metabolism and Fe-S cluster pathways, perhaps indicating that overexpression of the heterologous SOD, under conditions in which it gets primarily metalated with iron [23,27] affects iron homeostasis and thereby iron-sulfur cluster synthesis.

Both of the endogenous *E. coli* SodFMs, *Ec*SodA and *Ec*SodB, were detected in the WT cells but not from Δ*sodA*Δ*sodB* cells (Table 2). *Ec*SodA showed similar abundance in both wild type samples, whereas the abundance of *Ec*SodB was increased after induction of the heterologous protein. Also as expected, the *S. aureus* SodM was detected in all samples carrying the *Sa*SodM vector, consistent with leaky expression from pET plasmids, even in the absence of added inducer [64], but its abundance was dramatically increased (6-fold in wild type and 8-fold in Δ*sodA*Δ*sodB*) after induction (Table 2).

**Table 2:** Detection of superoxide dismutase isozymes in proteomic analyses of WT and Δ*sodA*Δ*sodB E. coli* cells.

| Protein | LFQ intensity average log<sub>2</sub> |  | Fold change after induction |  |
| --- | --- | --- | --- | --- |
| | WT | $2\Delta$ | WT | $2\Delta$ |
| *S. aureus* SodM | 34.26 | 34.22 | 6.39 | 8.68 |
| *E. coli* SodB | 30.89 | ND | 2.13 | ND |
| *E. coli* SodA | 29.28 | ND | 1.17 | ND |

Detection and abundance of endogenous *E. coli* SodA (MnSOD) and SodB (FeSOD) isozymes or heterologously expressed *S. aureus* SodM in the proteomic samples. ND = not detected in this sample.

Taken together, the proteomic data indicate that the Δ*sodA*Δ*sodB* strain shows no obvious physiological problems when transformed with an expression construct encoding a heterologous SOD isozyme that is expressed inside *E. coli* BL21 cells, even in the absence of inducer. Further, the data demonstrate that the Δ*sodA*Δ*sodB* strain is as efficient a host for production of a recombinant SodFM isozyme as the WT BL21 parent strain.

### The E. coli BL21 ΔsodAΔsodB strain is an excellent host for expression of diverse SodFMs

We have used the *E. coli* BL21 Δ*sodA*Δ*sodB* strain for production of recombinant SodFMs for *in vitro* characterisation. Our goal has been to characterise WT and mutated variants of SodFM isozymes to test their catalytic metal-preference [24,26]. We have used this approach to determine how mutations within the metal ion’s secondary coordination sphere influence metal preference [23], and how this biochemical parameter has evolved within the SodFM superfamily [21]. To characterise a SodFM’s catalytic metal-preference, we must accurately quantify its activity with both manganese and with iron, but for highly metal-preferring isozymes, one or other of these activities can be extremely low [21]. This *E. coli* strain was created to prevent contamination of recombinant SodFM preparations by the native *E. coli* SodFMs, SodA or SodB. The *E. coli* SodFMs exhibit highly metal specific (for manganese and iron, respectively) activity with very high catalytic efficiency [30,65], thus even low levels of contamination by them can interfere with such measurements.

We sought to validate that the *E. coli* BL21 Δ*sodA*Δ*sodB* strain makes a suitable expression host chassis for the synthesis of heterologous SodFM isozymes in forms loaded specifically with each target metal cofactor, in high abundance, free of contamination by the *E. coli* SodFMs, enabling measurement of their catalytic metal-preferences. To achieve this, we took advantage of our ability to perform in-gel assays. SOD enzyme activity can be assayed easily and semi-quantitatively in native polyacrylamide gel electrophoresis (PAGE). After resolving, the gels are exposed to a source of superoxide (light-dependent production catalysed by riboflavin) and a colorimetric dye (nitroblue tetrazolium, NBT) that develops colour in the presence of superoxide, forming a dark purple precipitate whose absence in regions of the gel indicates the presence of superoxide dismutase enzyme. We used this assay to demonstrate that the three bands of SOD activity (representing the two homodimeric, plus a heterodimeric form), which are readily detected in extracts from wild type *E. coli* BL21 in NBT-stained gels, were absent in equivalent extracts prepared from the Δ*sodA*Δ*sodB* strain (Fig. 1C). This confirmed that both SodA and SodB activity were eliminated from this strain, as expected.

Next, we compared the expression in the *E. coli* BL21 WT and Δ*sodA*Δ*sodB* strain of isozymes from each of the five SodFM subfamilies, which we previously defined by bioinformatic analyses [21], denoted SodFM1-5. These isozymes were the SodFM1 from *Staphylococcus aureus* (SodA), the SodFM2 from *Neisseria gonorrhoeae*, the SodFM3 from *Mycobacterium abscessus*, the SodFM4 from CPR *Parkubacteria* and the mitochondrial SodFM5 from *Homo sapiens*. Expression constructs for each isozyme, constructed in a prior study [21], were transformed into the BL21 WT and Δ*sodA*Δ*sodB* strain. Transformants were cultured in an M9 medium supplemented with 1 μM iron to mid-log phase, and then expression of the heterologous SodFM was induced by addition of 0.1 mM IPTG. To achieve selective metal-loading inside the bacterial cells, either 200 μM MnCl_2_ or 100 μM FeSO_4_ was added to the growth medium at the point of induction. Cells were harvested after 4 h incubation and cell lysates analysed by gel electrophoresis to assess expression and metal-loading.

Initial SDS-PAGE analysis showed that all SodFM isozymes, with the exception of the SodFM3 from *Mycobacterium abscessus*, was successfully overexpressed inside both WT and mutant host strains of *E. coli* (Fig. 4). Notably, when an increased quantity of biomass (50 μg) was loaded on SDS-PAGE, even the SodFM3’s expression could be observed, albeit at relatively low levels (Supp. Fig. S4). In a prior study, expression of this SodFM3 was improved by expression within *E. coli* cells by an additional plasmid that provides additional expression of tRNAs for rare codons [21]. Thus, all five of the SodFM isozymes were expressed in the BL21 Δ*sodA*Δ*sodB* host strain. In all cases, the expression level obtained with the BL21 Δ*sodA*Δ*sodB* strain was analogous to that obtained in the parental wild type BL21 strains (Fig. 4; Supp. Fig. S4).

**Figure 4:**
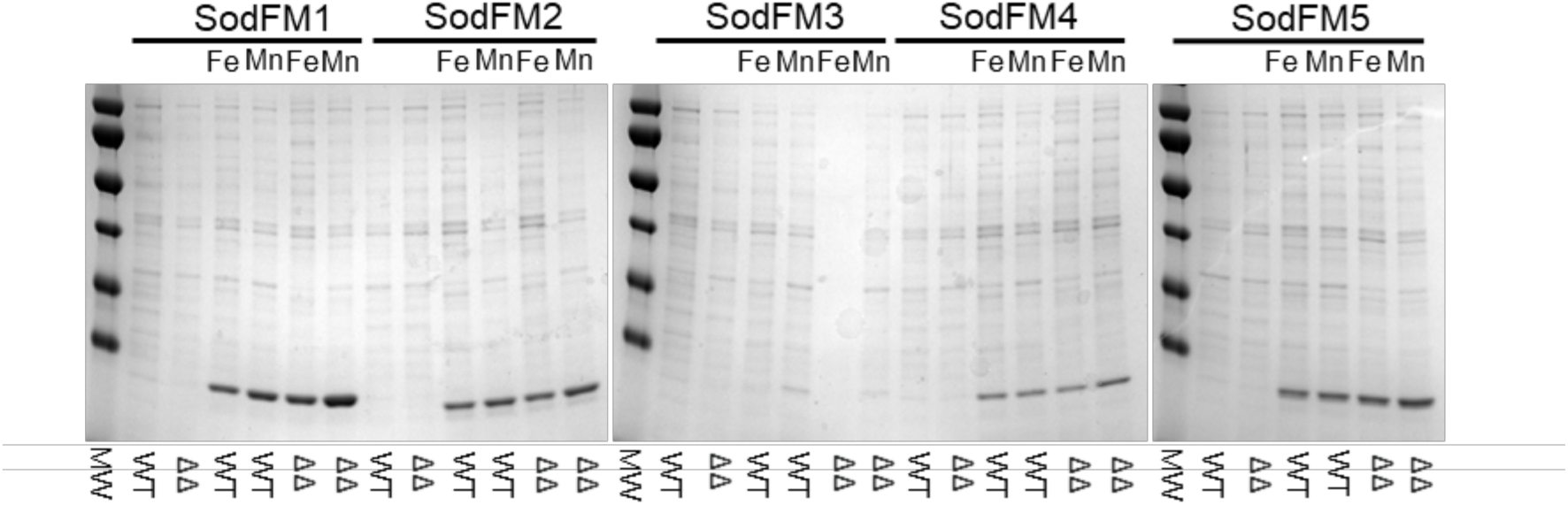
SDS-PAGE analysis of lysates from *E. coli* expressing diverse SodFMs. The BL21 wild type (WT) or Δ*sodA*Δ*sodB* (ΔΔ) strains were each transformed with pET22b constructs for the heterologous expression of the SodFM1 from *Staphylococcus aureus* (*Sa*SodA), the SodFM2 from *Neisseria gonorrhoeae*, the SodFM3 from *Mycobacterium abscessus*, the SodFM4 from CPR *Parkubacteria* or the SodFM5 from *Homo sapiens*. Each strain was cultured to mid-log phase in M9 minimal medium containing glucose and supplemented with 1 µM FeSO_4_. Expression was induced by addition of 100 µM IPTG, at which point selected cultures were also supplemented with either 200 µM FeSO_4_ (Fe) or 200 µM MnCl_2_ (Mn), and cultures were incubated at 37°C overnight. Cells were harvested, washed and lysed in 20 mM Tris, pH 7.5, 100 mM NaCl through multiple (x3) freeze-thaws using liquid N_2_. Equal amounts (1 µg) of each extract were resolved by SDS-PAGE on a 15% acrylamide gel alongside molecular weight markers (MW) and stained with Coomassie Brilliant Blue.

To test the activity of the overexpressed enzymes, we resolved equal amounts of each crude soluble extract (3 μg) on native PAGE and negatively stained the resulting gel with NBT/riboflavin (Fig. 5A). In each case, an abundant band of activity was clearly detected in both the BL21 wild type and the Δ*sodA*Δ*sodB* strain for all SodFM isozymes, including the SodFM3 despite its relatively low expression. In the samples from both *E. coli* strains, the SodFM1, SodFM3, SodFM4 and SodFM5 isozymes showed much more activity in the samples cultured in the presence of manganese than those cultured in the presence of iron, whereas the opposite was the case for the SodFM2, consistent with the metal-preferences of the five tested SodFMs that were previously demonstrated [21]. Importantly, lysates prepared from WT cells exhibited additional bands of activity that were absent in the crude lysates prepared from the BL21 Δ*sodA*Δ*sodB* strain (Fig. 5A).

**Figure 5:**
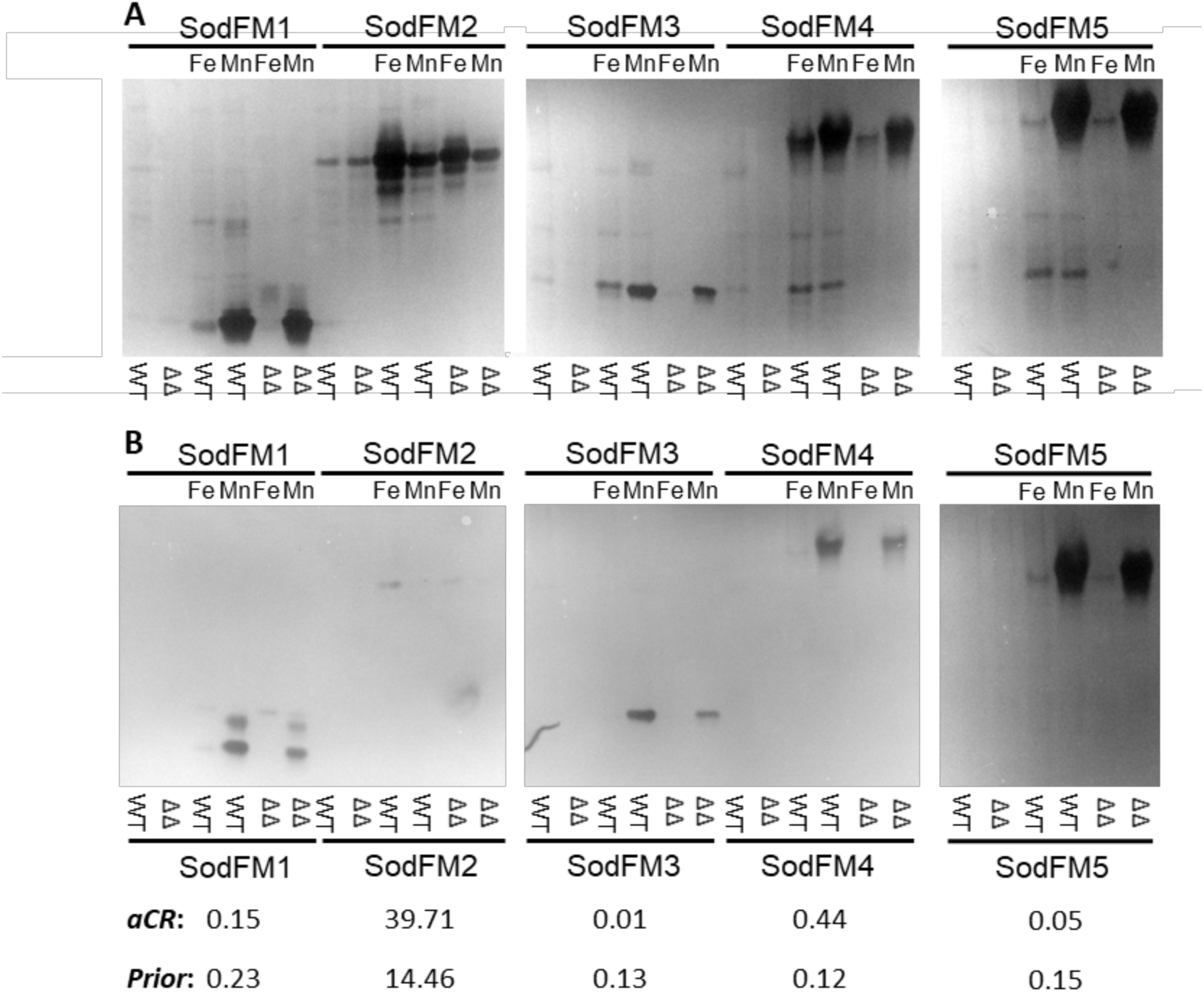
In-gel SOD activity assay of lysates from *E. coli* expressing diverse SodFMs. The same cell extracts, prepared from the BL21 wild type (WT) or Δ*sodA*Δ*sodB* (ΔΔ) strains transformed with the SodFM expression constructs, and selectively loaded with Mn or Fe, as described in Fig. 4, were analysed in-gel for enzymatic activity. Equal amounts (3 µg) of each extract were resolved by native PAGE on pairs of 15% (v/v) acrylamide gels at pH 8.8. One gel (**A**) was incubated in water alone, and the other gel (**B**) was incubated in a solution of 0.3% (v/v) H_2_O_2_ at 4°C, for 30 min and then both gels underwent negative staining with NBT/riboflavin to detect regions of superoxide dismutase activity. Gels were scanned and inverted for clarity. Densitometry was used to quantify activity in each gel and used to calculate the approximate cambialism ratio (aCR) [23], shown below the gels along with the equivalent result from a prior study [21].

Hydrogen peroxide is a specific inhibitor of the iron-loaded forms of SodFMs but has negligible effects on manganese-dependent SodFM activity [21]. To estimate metal-loading in crude extracts, replicate native gels were soaked in 0.3% (v/v) H_2_O_2_ prior to negative staining for enzymatic activity (Fig. 5B). As expected, when gels stained with or without prior peroxide treatment were compared (Fig. 5), all bands of activity in lysates prepared from cells cultured in the presence of iron showed significant inhibition of activity, whereas those from cells cultured in the presence of manganese showed little inhibitory effects. This indicated that the over-expressed protein had, in each case, primarily been loaded with the metal ion that was added in excess to the bacterial culture medium at the point of induction. Quantification of the activity bands in the gels through densitometry enabled the assessment of the relative activity with iron and with manganese for each SodFM isozyme and calculation of the approximate cambialism ratio (aCR; cambialism ratio is defined as the measured Fe-dependent activity divided by the manganese-dependent activity) using a published protocol [21]. This analysis demonstrated that the metal-preference of these five SodFM isozymes was in line with those previously reported (Fig. 5).

In-gel analyses are insufficiently sensitive to assess the levels of cross-metal contamination of the heterologously expressed SodFMs, especially for those that exhibit highly metal-preferring catalysis. To confirm quantitatively the metal-loading of the recombinant SodFMS during expression inside cells of the *E. coli* BL21 Δ*sodA*Δ*sodB* strain, we purified samples of the *S. aureus* SodA (*Sa*SodA) protein after expression during growth in media containing either excess manganese or iron. Each sample of protein was purified according to existing protocols we have used to generate samples for *in vitro* studies [27]. Triplicate samples were analysed for metal content using ICP-OES. The data demonstrated highly specific metal-loading of *Sa*SodA was achieved during over-expression inside the *E. coli* cells when cultured under these conditions (Fig. 6).

**Figure 6:**
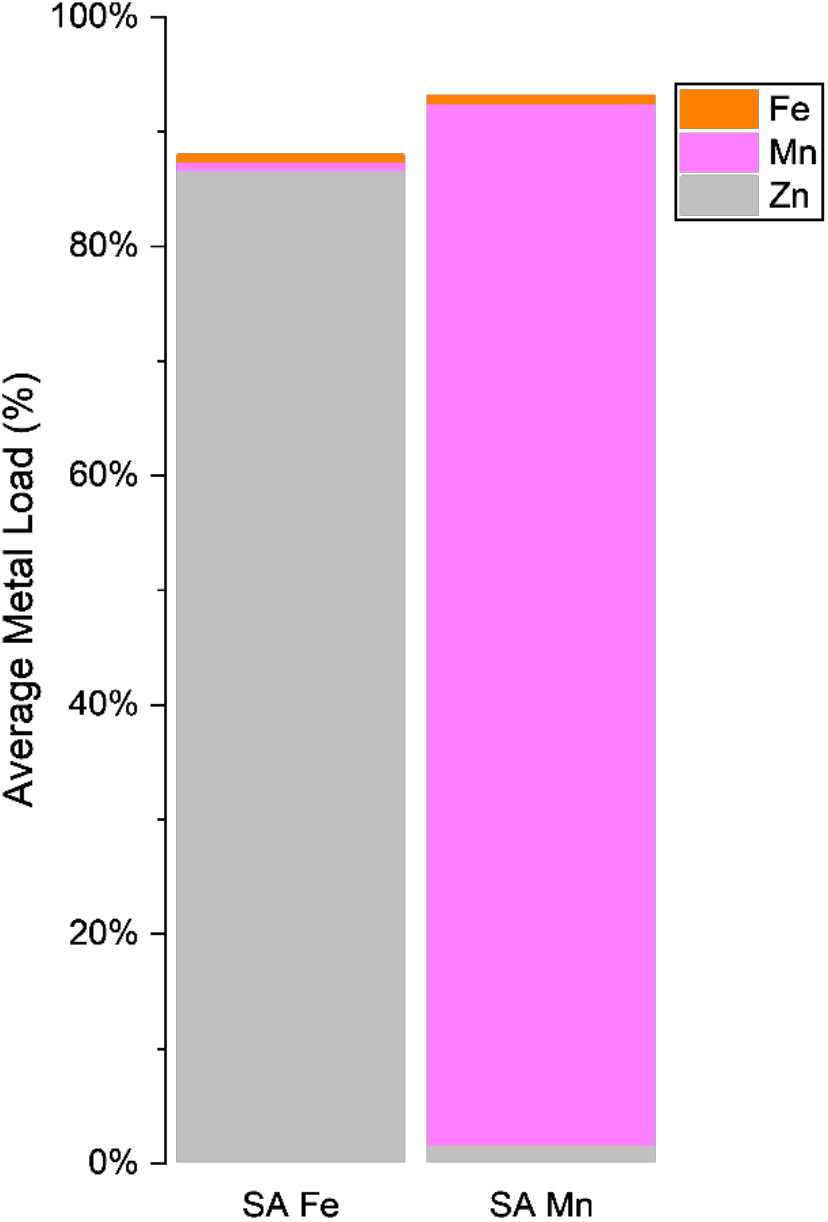
Selective metal-loading of heterologous, over-expressed SodFMs is possible in the *E. coli* Δ*sodA*Δ*sodB* strain. Recombinant *Sa*SodA from *S. aureus* was over-expressed inside cells of the *E. coli* BL21 (DE3) Δ*sodA*Δ*sodB* strain. Cells were cultured in M9 medium, and then at the point of protein induction with IPTG, cultures were supplemented with either 100 µM FeSO_4_ (left) or with 200 µM MnCl_2_ (right) to achieve loading of the recombinant proteins with these respective metals. The over-expressed proteins were purified by standard methods [23,27,68], and the loading of each recombinant isoform with iron, manganese or zinc was measured by ICP-OES, with protein quantitation in the ICP samples determined using their S content.

Taken together, these results demonstrated that the *E. coli* BL21 Δ*sodA*Δ*sodB* strain is an ideal expression chassis for production of recombinant SodFM isozymes, from all five identified phylogenetic subfamily lineages, and is capable of producing them in a form that is largely loaded with a specific metal cofactor, thereby enabling in vitro assessment of its metal-preference of catalysis.

## Discussion

We previously constructed a strain of *E. coli* BL21 for the purpose of creating a microbial cell factory for the efficient synthesis of recombinant heterologous superoxide dismutase enzymes free of contamination by its endogenous cytosolic SodFM isozymes. The native *E. coli* SodFMs exhibit very high metal-preference of their catalysis, with *Ec*SodA being active only with manganese and *Ec*SodB only with iron, each with very high catalytic turnover [30,65]. Therefore, even low levels of contamination by the *E. coli* SodFMs can contribute to the measured SOD activity of recombinant protein preparations, especially when biochemically characterising an isozyme with low catalytic turnover (for example, when measuring turnover from a manganese-preferring isozyme when wrongly loaded with iron and *vice versa*). Additionally, while the expression of the iron-dependent *E. coli* SOD, *Ec*SodB, is largely constitutive, the manganese-dependent *E. coli* SOD, *Ec*SodA, is induced under stress conditions [66]. Furthermore, it is feasible that the endogenous *E. coli* SODs might also assemble into heterodimeric forms with the heterologous SodFM, which would result in contamination even after purification using a protein-tagging strategy. Thus, it is unavoidably unclear to what extent each of these isozymes will contribute through their contamination to a given extract, interfering not only with the measurement of catalytic activity but also to elemental analysis to determine the metal content of recombinant protein preparations of heterologously expressed SodFMs. Our *E. coli* BL21 Δ*sodA*Δ*sodB* strain was designed to overcome this contamination problem, enabling us to produce large quantities of the purified recombinant enzymes for detailed biochemical and biophysical characterisation [21,22]. Use of this strain enables the production of contaminant-free recombinant SodFMs without the use of protein tags to facilitate the purification process. It also created a system for screening the activity of SodFMs in a higher throughput manner, avoiding the need for full purification of target isozymes in each metal form by assaying the recombinant enzyme direct from crude bacterial lysates, either in solution or in gel activity assays. The latter provided us with a tool that enabled us to survey a diverse set of 60 SodFM isozymes from across the entire phylogenetic tree, as well as more than 100 different mutated variants of those isozymes, to determine the evolutionary history of this widely distributed metalloenzyme family [21].

In this study, we sought to characterise this mutant strain of *E. coli* BL21 to identify any physiological defects which it might exhibit and to validate it as a new tool for use in the study of the remarkable SodFM family of metalloenzymes. Based on prior observations made with a mutated strain of Type A *E. coli*, strain K-12, we anticipated that we would observe physiological defects in our *E. coli* BL21 Δ*sodA*Δ*sodB* strain. In particular, it was shown that the *E. coli* K-12 Δ*sodA*Δ*sodB* strain failed to grow in a minimal medium supplemented with glucose as its sole carbon source [39], a phenotype proposed to result from superoxide-mediated damage to Fe-S clusters in essential enzymes of carbon metabolism [67]. We observed similar phenotypes for our *E. coli* BL21 Δ*sodA*Δ*sodB* strain. Notably, we found no significant changes in the genome sequence of the strain, ruling out a strong selection pressure giving rise to suppressor mutations during the selection stages of genetic modification. Finally, we observed only minor differences in the expression of the proteome between WT and Δ*sodA*Δ*sodB* cells in LB medium, further confirming that the physiology of the Δ*sodA*Δ*sodB* strain presents no problems for its use as a cell factory for production of recombinant SodFMs.

We demonstrated the usefulness of this *E. coli* BL21 (DE3) Δ*sodA*Δ*sodB* strain for production of recombinant SodFM enzymes. First and foremost, the strain can produce large quantities of heterologous isozymes from across the SodFM phylogenetic tree, assuring that preparations are free of contamination by *Ec*SodA and *Ec*SodB. The resulting samples can be readily purified to near-homogeneity by liquid chromatography, without need for affinity tags, resulting in samples that free of contamination by *E. coli* SODs, which are suited for biochemical characterisation or structural determination [21–24]. Metal-loading control can be achieved to a significant extent by culturing the strain under carefully defined metal conditions in M9 minimal medium. Unfortunately, some mixed metal loading of heterologous proteins are unavoidable in-cell: both of the contaminating metals, Fe and Zn, are needed for growth so cannot be omitted, and therefore are also present in low-levels in the purified preparations. Nonetheless, the strain can make protein predominantly loaded with a target metal ion. We have optimised these conditions specifically for our extensive studies of the SodFM1 isozymes from *S. aureus*, and these conditions produce highly selective metal loading with either manganese or iron based on supplementation of the growth medium with these metals. Importantly, the strain also enables a higher throughput assay that enables the assessment of relative activity of isozymes with manganese and iron as cofactor, using an in-gel activity assay combined with the use of the selective inhibitor of iron-dependent SodFM activity, hydrogen peroxide [21]. The combination of this *E. coli* strain, a microbial cell factory that avoids contaminating contributions to detected activity from endogenous SodFMs, with the convenient in-gel assay and the conserved selective inhibition by peroxide, enables assessment of the metal-preference of diverse SodFMs without the need for time-consuming and costly purification of two metal forms of each isozymes under study.

## Conclusions

Here, we have characterised the physiology and utility of an *E. coli* BL21 Δ*sodA*Δ*sodB* strain for production of recombinant SodFM proteins for *in vitro* studies. Genomic analysis verified the targeted gene deletions, without significant off-target effects. Growth, expression, elemental analysis, and proteomic data confirmed a lack of physiological defects of the mutant strain, except for a known inability to grow on glucose, which we have demonstrated can be overcome by heterologous expression of a SodFM isozyme. We demonstrate the strain can efficiently produce diverse recombinant SodFMs, including precise control of metal-loading of the heterologously expressed protein. The *E. coli* strain is thus a useful microbial cell factory for production of recombinant SodFMs, which should find widespread utility as expression host of choise, enabling improved production of protein for studies of the biochemical, biophysical and structural properties of this remarkable family of metalloenzymes.

## List of Abbreviations

aCR: approximate cambialism ratio
CR: cambialism ratio
IMAC: immobilised metal affinity chromatography
IPTG: isopropyl β-D-1-thiogalactopyranoside
LB medium: Luria-Bertani medium NBT: nitroblue tetrazolium
PAGE: polyacrylamide gel electrophoresis
SDS-PAGE: sodium dodecyl sulfate polyacrylamide gel electrophoresis
SNP: single nucleotide polymorphism
SOD: superoxide dismutase
SodFM: iron- or manganese-dependent superoxide dismutase

## Declarations

### Ethics approval and consent to participate

Not applicable.

### Consent for publication

Not applicable.

### Availability of data and materials

The genomic sequencing data supporting the conclusions of this article are available in the BioProject repository (BioProject ID: PRJNA1272326; http://www.ncbi.nlm.nih.gov/bioproject/1272326). The proteomic data supporting the conclusions of this article are available in the PRIDE repository. All other data supporting the conclusions of this article are included within the article and its additional file(s), and source data is available from the corresponding author on reasonable request.

### Authors’ contributions

The preparation of DNA and protein samples for sequencing and proteomics were performed by AC under supervision by RM and ME. Growth analyses were performed by DG and gene expression analyses were performed by JP. The expression, analysis and purification of SodFMs were performed by RM, with assistance from ME. Genomic data analysis was performed by JG. Mass spectrometry analyses were performed by AM and AS, and proteomic data were analysed by RM with assistance from ME, AC, AM and AS. The project was conceived by ME, TEKF and KJW and supervised and managed by KJW. The manuscript was written by KJW with contributions from all other authors. All authors read and approved the final manuscript.

