## Supplementary information for "An engineered biofactory for efficient production of diverse recombinant superoxide dismutase isozymes loaded with specific metal ions for biochemical characterization"

**Short title:** Genomic and proteomic analysis of *Escherichia coli* BL21 Δ*sodA*Δ*sodB* cell factory

**Keywords:** *Escherichia coli*; superoxide dismutase; cell factory; proteomics; genome

### Supplementary information

### Supplementary Tables

**Supp. Table S1: Sequences of oligonucleotide primers used in this study.**

| Gene name | Gene target | Primer | Sequence |
|---|---|---|---|
| *fecA* | Ferric citrate transporter | F | ACTCCAACCAGACCAACGACAC |
| | | R | ATCGTAACGTGCCTGCGTTTCC |
| *mntH* | $Mn^{2+}/Fe^{2+}:H^{+}$ symporter MntH | F | ACCGACCTGGCGGAATTTATTGG |
| | | R | CCGCACCTTGCAACAACGAAAC |
| *feoA* | Ferrous iron transporter, protein A | F | CGTGAAATCAGCCCGGCATATC |
| | | R | AGGAGCCAGGTAACATGCCAAG |
| *rrsA* | 16S ribosomal RNA | F | CTCTTGCCATCGGATGTGCCCA |
| | | R | CCAGTGTGGCTGGTCATCCTCTCA |
| *hcaT* | 3-phenylpropionic transporter | F | GCTGCTCGGCTTTCTCATCC |
| | | R | CCAACCACGCTGACCAACC |

### Supplementary Figures

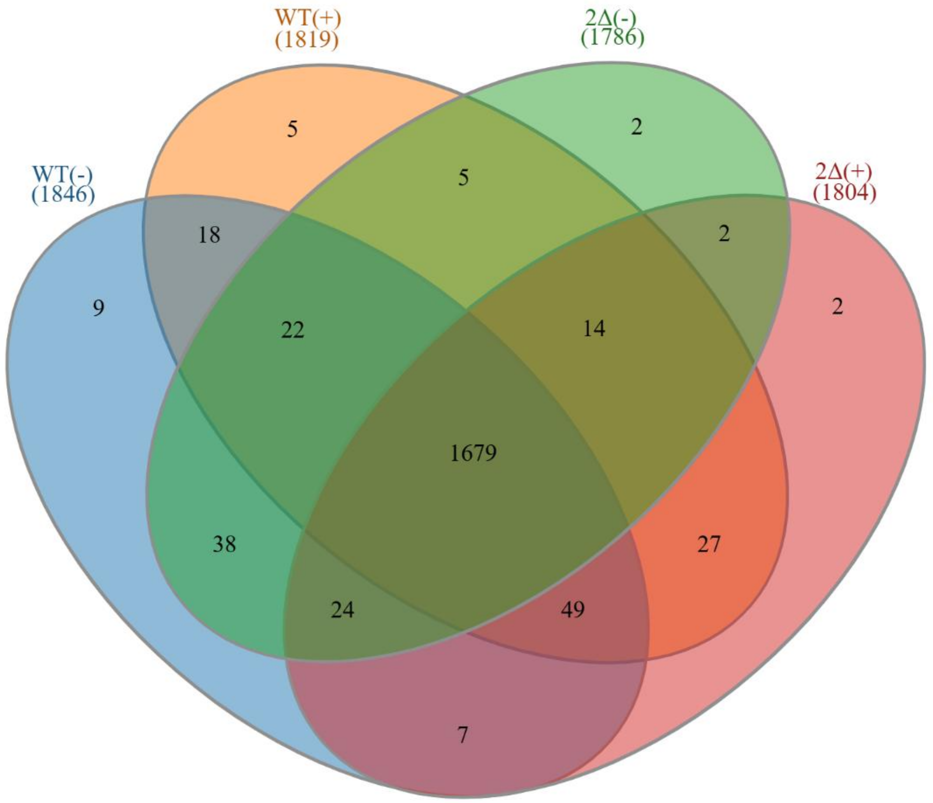


**Supplementary Figure S1: Comparison of the proteins detected in each sample.**

Venn diagram illustrating the detection of proteins in each proteomic analyses of *Escherichia coli* BL21 (DE3) (WT) or the Δ*sodA*Δ*sodB* strain (ΔΔ). Cell were cultured either in the absence (-) or the presence (+) of the inducer IPTG to induce the expression of the heterologous protein SodM from *Staphylococcus aureus*. A total of 1,679 proteins detected were common to all four samples.

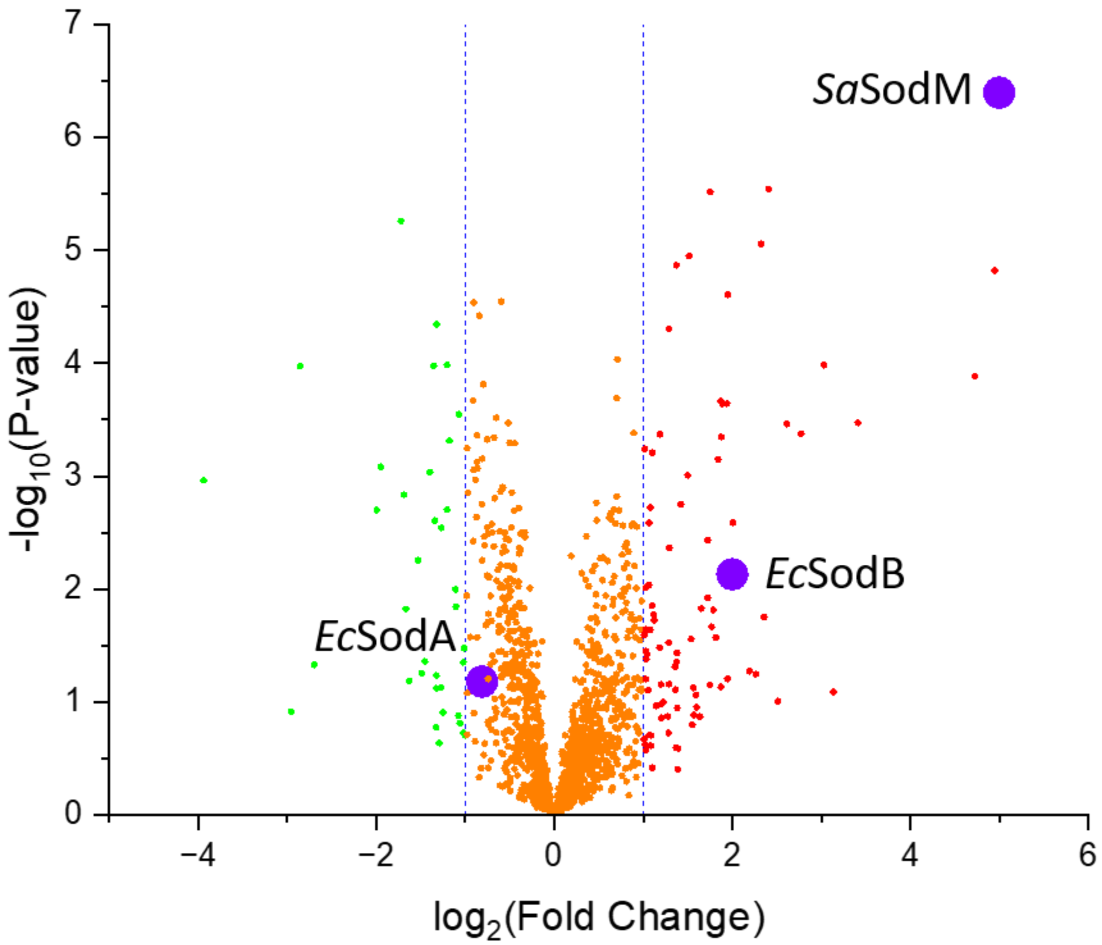


**Supplementary Figure S2: Proteomic comparison of protein abundances in wild type *E. coli* BL21 (DE3) cells before and after induction of expression of *S. aureus* SodM.**

Proteins were extracted from aliquots (1 mL) of cultured *E. coli* BL21 (DE3) wild type carrying the pET22b-SodM construct, with and without induction (100 μM IPTG) of expression of the heterologous SodM protein. Cells were lysed in SDS buffer by sonication, equal amounts of protein (35 μg) digested with trypsin, and resulting peptides were purified. Peptide mixtures (2 μg) analysed by LC-MS. Peptides were assigned to proteins with MaxQuant using an *E. coli* reference database, and then changes in protein abundance analysed with Perseus. The resulting volcano plot compares the WT strain without added IPTG inducer to that with induction of the heterologous protein. The red circles represent proteins that were up-regulated (>1 $log_2$), the green circles represent the proteins that were down-regulated (<1 $log_2$), and all proteins found to show no change in abundance are represented by orange circles. The large blue circles represent the heterologously expressed *S. aureus* SodM protein and the endogenous SodA and SodB enzymes found in *E. coli*. The dotted blue lines represent the log2 fold-change cutoff used.

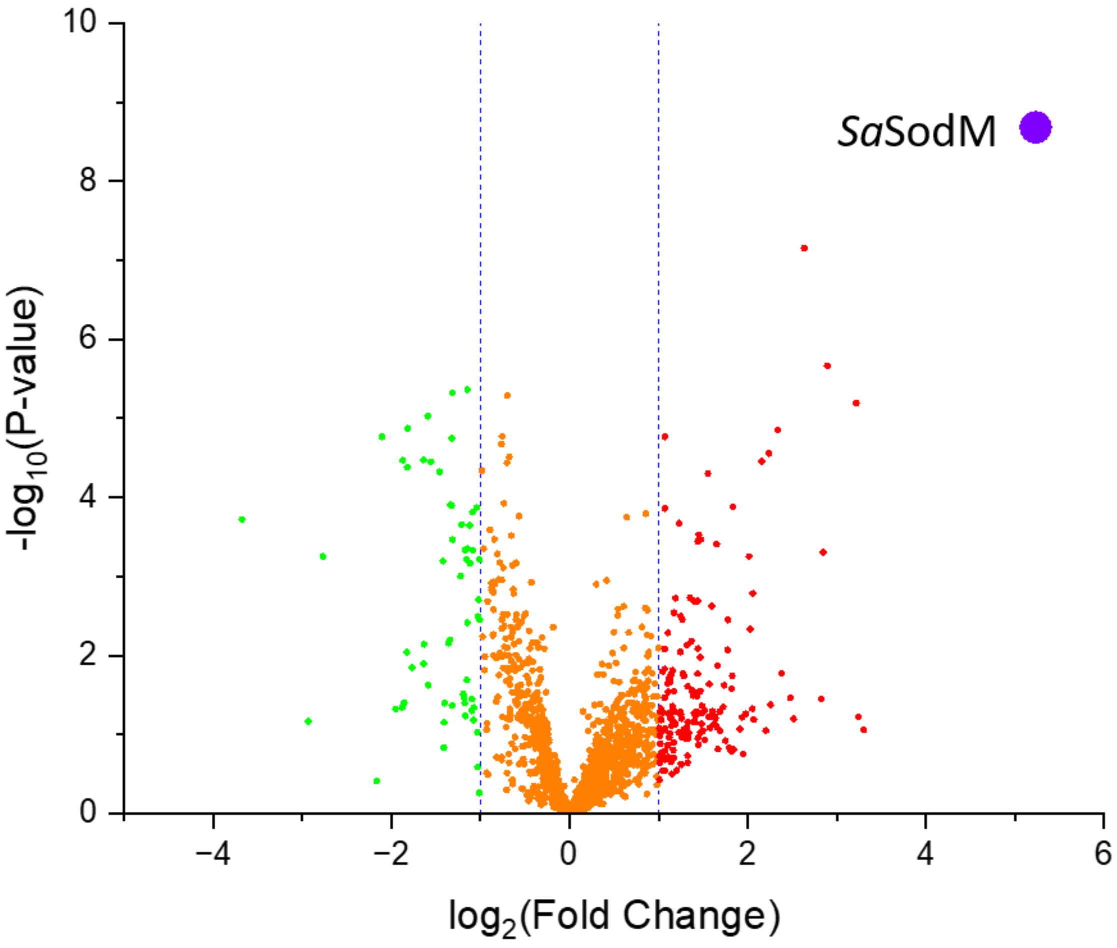


**Supplementary Figure S3: Proteomic comparison of protein abundances in *E. coli* BL21 (DE3) Δ*sodA*Δ*sodB* cells before and after induction of expression of *S. aureus* SodM.**

Proteins were extracted from aliquots (1 mL) of cultured *E. coli* BL21 (DE3) Δ*sodA*Δ*sodB* cells carrying the pET22b-SodM construct, with and without induction (100 μM IPTG) of expression of the heterologous SodM protein. Cells were lysed in SDS buffer by sonication, equal amounts of protein (35 μg) digested with trypsin, and resulting peptides were purified. Peptide mixtures (2 μg) analysed by LC-MS. Peptides were assigned to proteins with MaxQuant using an *E. coli* reference database, and then changes in protein abundance analysed with Perseus. The resulting volcano plot compares the Δ*sodA*Δ*sodB* strain without added IPTG inducer to that with induction of the heterologous protein. The red circles represent proteins that were up-regulated (>1 $\log_2$), the green circles represent the proteins that were down-regulated (<1 $\log_2$), and all proteins found to show no change in abundance are represented by orange circles. The large blue circle represents the heterologously expressed *S. aureus* SodM protein. The dotted blue lines represent the log2 fold-change cutoff used.

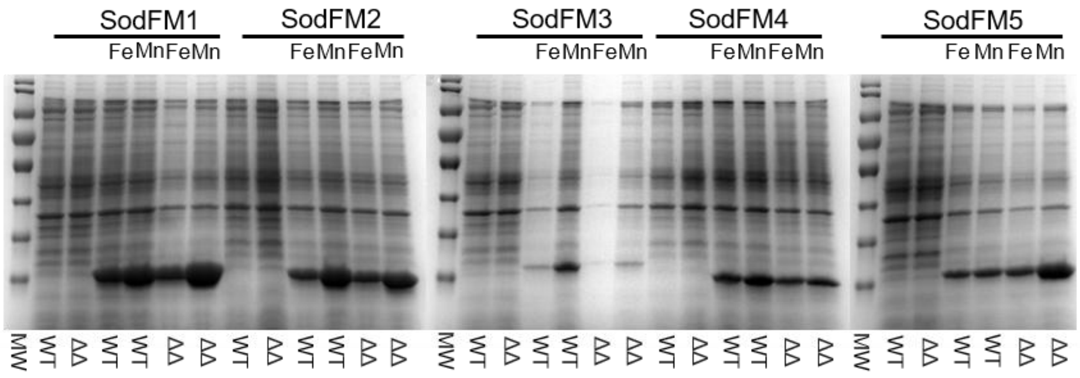


**Supplementary Figure S4: SDS-PAGE analysis of lysates from E. coli expressing diverse SodFMs.**

The BL21 wild type (WT) or Δ*sodA*Δ*sodB* (ΔΔ) strains were each transformed with pET22b constructs for the heterologous expression of the SodFM1 from *Staphylococcus aureus* (SodA), the SodFM2 from *Neisseria gonorrhoeae*, the SodFM3 from *Mycobacterium abscessus*, the SodFM4 from CPR *Parkubacteria* or the SodFM5 from *Homo sapiens* [1]. Each strain was cultured to mid-log phase in M9 minimal medium containing glucose and supplemented with 1 µM $FeSO_4$. Expression was induced by addition of 100 µM IPTG, at which point the cultures were also supplemented with either 200 µM $FeSO_4$ (Fe) or 200 µM $MnCl_2$ (Mn) for selective metal-loading, and cultures were incubated at 37°C overnight. Cells were harvested, washed and lysed in 20 mM Tris, pH 7.5, 100 mM NaCl through multiple (x3) freeze-thaws using liquid $N_2$. Equal volumes (5 µL) of each extract were resolved by SDS-PAGE on a 15% (v/v) acrylamide gel alongside molecular weight markers (MW), and stained with Coomassie Brilliant Blue.
